# *Valeriana Officinalis* Crude Extracts Impair Hair Cell Regeneration in Larval Zebrafish Lateral Line

**DOI:** 10.1101/2019.12.30.891366

**Authors:** Roberto Rodríguez-Morales, Alexis Santana-Cruz, Tiffany Tossas-Deida, Aranza Torrado-Tapias, Martine Behra

**Affiliations:** School of Medicine, Department of Anatomy and Neurobiology, University of Puerto Rico, Medical Sciences Campus (UPR-MSC), San Juan, USA

**Keywords:** Hair cell, lateral line, regeneration, zebrafish, valproic acid, valerian, crude root extracts, hearing disorders, balance disorders

## Abstract

Irreversible hair cell (HC) loss in the inner ear is the leading cause for hearing and balance disorders. Discovery of therapeutic molecules preventing HC death and promoting regeneration, which does not occur in mammals like it does in lower vertebrates, is of major interest. In fish, HCs are also found in a superficial mechano-sensory organ called the lateral line (LL). LL-HCs are exposed to surrounding waters and are accessible to waterborne molecules providing a potent mean to study *in vivo* HC stability and regeneration. Commercial small molecule libraries were tested in screens for HC survival and regeneration in zebrafish, but ethnobotanical pharmacopeias remain totally unexplored because of the challenge that such complex mixtures represent. A rapid and cost-effective first-pass assay informing about the regenerative potential of an extract is therefore critical before embarking on cumbersome purification steps. We chose to test Valerian crude root extracts (Val), which are typically composed of more than 150 different components, amongst which is a main and abundant compound: valeric acid (VA). VA discovery and purification led to the commercialization of a synthetic analog: Valproic acid (VPA) which is a first-line drug for epilepsy and bipolar disorders that was also shown to significantly hamper LL-HC regeneration. We reasoned that if Val is not toxic, it would elicit effects like VPA. Thus, we synchronously ablated HCs in 5-day post-fertilization (dpf) larvae and monitored regeneration over the following 3 days in the presence of Val at the highest well-tolerated concentration (Val_1_ = 1mg/ml), or VPA (= 150μM) as previously published. Both treatments significantly decreased HC regeneration without affecting HC-survival suggesting a similar mode of action. Furthermore, Val application as early as 3dpf and prolonged for up to 4 days did not affect larval survival, indicating that reduced HC-regeneration was not due to overall toxicity. Taken together, Val and VPA-treatments displayed a comparable response in a simple and up-scalable HC-regeneration assay which is an in-first-pass potent approach for drug discovery.

## Introduction

Hearing disabilities are affecting close to 7% of the world population creating a major burden on global human health (1). About 90% of hearing impairments are sensorineural hearing losses, meaning that they are affecting the hair cells (HCs) in the sensory epithelium of the inner ear (= organ of Corti in the cochlea), or their innervation via the auditory nerve (2). Genetic and environmental factors cause sensorineural hearing loss which can be categorized accordingly, in genetic forms of hearing loss (GFHL), noise-induced hearing loss (NIHL), drug-induced hearing loss (DIHL) and age-related hearing loss (ARHL) (for review (3)). Whatever the cause of HC loss is, it is irreversible in mammals who strikingly miss the regenerative power found in lower vertebrates (for review (4)). Originally thought to be limited to cold blooded animals (5, 6), HC regeneration was also demonstrated in birds (5, 6) and could be partially induced in murine cochlea (7, 8). In addition, vestibular HCs of the inner ear which are gravity/balance sensors and gaze stabilizers, can regenerate to some extent in adult mammals (9), arguing that HC regeneration could be restored in humans. To this effect, a range of strategies are pursued such as molecular therapies to counteract drugs or noise ototoxicity, cell- and gene-therapies to compensate mutation in genes causing HC loss and to induce HC regeneration (for review (10)).

Another major line of research is drug discovery in search of molecules able to induce or remove the cellular brakes preventing HC regeneration. In this domain, the genetic tractable zebrafish has proven to be a valuable animal model because it also has regenerating HCs in a mechano-sensory organ called the lateral line (LL) (11). This superficial structure is constituted of stereotypically distributed superficial sensory patches called neuromasts (NMs), which are comprised of central HCs surrounded by support cells (SCs). The LL provides an easy experimental access to interfere with HC-development or regeneration which can be monitored *in vivo* after addition of waterborne compounds (12). Rapid development of unlimited number of small larvae make it optimal for high-throughput screens (13). Search of protective molecules against ototoxicity (14) and assessment of ototoxicity of anti-cancer drugs were performed (15, 16). Screens of FDA-approved small molecule libraries revealed several otoprotective compounds that were further validated in mice (17) and a few modulators of HC regeneration (18), thus clearly validating these approaches.

However, screens so far have only interrogated single purified compounds leaving a major resource untapped: natural products (NPs). NPs can be broadly defined as plant, animal or fungi extracts with different degrees of purification for which effects have been properly documented in the abundant ethnobotanical literature (19). Systematic testing of NPs is daunting for many reasons and not the least being that reports on active preparations are challenging to reproduce. The mode of preparation is often highly variable, the crop origin is non-descript, the harvesting methods are unknown, and all can greatly affect the composition of any given extract. Notably, purification steps can be expensive, labor-intensive, and time-consuming, making it unpractical to embark on such steps without further knowledge of the actual potential of a crude extract.

To address this gap, we explored the possibility to exploit a cost-effective first-pass assay for testing unknown complex mixtures for their potential to interfere with HC-regeneration. We chose a simple HC regeneration assay that we (20) and others (21) have previously performed. We triggered massive and synchronous HC loss with a punctual copper treatment (Cu^++^, 10μM for 2h) (22) and then followed HC regeneration over the next 3 days, which is sufficient time for restoring NMs to the pretreatment state (23–26). As proof of principle, we tested crude root extracts of *Valeriana officinalis*, commonly called Valerian (Val). Similar preparations typically contained more than 150 different ingredients (27), but with one main component being always present: valeric acid (VA). VA led to the synthesis of a powerful chemical analog named valproate (or valproic acid, VPA) which is the top-prescribed psychoactive drug worldwide (28). More relevant here, VPA was also shown to severely reduce LL-HC regeneration in zebrafish larvae (29, 30). Our rationale was that if Val is not toxic, it would contain enough VA to elicit comparable effects to purified synthetic VPA, and therefore demonstrate that a non-purified active compound when present in a complex mixture can be detected with the proposed screening assay.

Consequently, after ablating all HCs in 5dpf larvae, we treated them with 150μM VPA (VPA_150_) as previously published (29, 30), or with the highest well-tolerated concentration of Val (Val_1_=1mg/ml) for the following 3 days. We found a significant decrease in HC-regeneration with both treatments which was not due to HC toxicity of neither treatment, suggesting a similar mode of action. To exclude that the reduced number of regenerating HCs was due to compromised larval health and/or survival, we treated animals from 3dpf and up to 7dpf with Val_1_ or VPA_150_ and found no significant deleterious health effect or mortality, arguing that reduced HC-regeneration was specific and not due to general toxicity. Taken together, we showed that the potential of a complex mixture of unknown composition to interfere with HC regeneration can be detected in a simple HC regeneration screen. Such a rapid, first-pass, up-scalable approach can be used before embarking on further purification steps to narrow down on the active compound in a NP. Thus, our work is proposing a new avenue to tap into unexploited ethnobotanical pharmacopeias in the search of therapeutic molecules capable to impair or promote HC regeneration.

## Materials and Methods

### Fish Husbandry and Care

We raised wild type (TUB × AB = TAB-5) and transgenic Tg (*tnks1bp1: EGFP*) adult zebrafish in system water (SW) with a stable pH = 7 and conductivity = 920-1100 micro-Siemens, in a semi-closed system (from Techniplast). Husbandry was performed as described (31) and following the National Institutes of Health guidelines for the care and use of laboratory animals and authorized by IACUC (protocol # A880110). We crossed couples for spawning and collected embryos in Petri dishes in blue water (BW = 20mL methylene blue, 0.54g ocean sea salt, 10L distilled water) and stored them in an incubator at 28°C. The following day, we replaced BW with SW and renewed it daily while raising embryos/larvae at 28°C to the desired stage.

### HC regeneration assay

We destroyed LL-HCs in 5dpf wild type larvae by exposing them for two hours to freshly prepared Copper Sulfate (II) solution (Cu^++^, Sigma # PHR1477-1G) at 10μM in Holtfreiter’s buffer 1X (HB-10 X stock = 30g NaCl, 2.6g CaCl_2_, 1g KCl, 1L RO water). Next, we washed larvae 6x 1min in HB-1X before dividing them in three groups raised further in one of the specified medium: (1) control (SW alone), (2) Val treated (SW + Val_1_= 1mg/ml), and (3) VPA treated (SW + VPA_150_ = 150μM). We renewed all media daily. To count the number of regenerated HCs at each desired stage (+ 0hour post Cu^++^ treatment (hpt), +24hpt, +48hpt and +72hpt), we collected the desired number of animals and treated them with FM1-43X (Invitrogen #F35355, 3μM /1min in the dark). Next, we anesthetized them with cold (or 0.002% MS-222) and mounted larvae in ice cold 4% methylcellulose on coverslips. We observed and imaged NMs (see below) on an inverted Zeiss microscope (Axiovert). After live observation we fully anesthetized larvae with ice-cold water (or 0.02% MS-222) and fixed them in fixing solution (FIX = 4% PFA + 4% sucrose) O/N at + 4°C. We washed the animals the next day in 6×10min PBS-1X + 0.1% Tween-20 (= PBST) and counterstained them with DAPI (10nM/20 min). Next, we dehydrated them in 100% glycerol and mounted them in Aqua Poly-mount ((PolySciences, #18606-20) for further imaging.

### Valeriana officinalis and VPA preparation and treatments

We prepared aqueous Valeriana plant extracts solutions from dry powdered Valerian roots, which were organically grown and minimally processed (Pacific Botanicals, Grants Pass, OR www.pacificbotanicals.com). We kept the dry powder at 4°C in a tightly sealed opaque container and prepared stock solution (Val= 100mg/mL) in SW and we diluted to the desired final concentration as needed in SW. We prepared stock solution for Valproic acid (VPA, Sigma-Aldrich #P4543) at 200mM in distilled water and diluted to the final desired concentration in SW as needed.

### NMs imaging and HCs counting

We imaged at least 3 head NMs (SO1, SO2 and SO3 or IO1, IO2 and IO3) and 3 trunk NMs (L1 to L5) and made Z-stack images (1μM thick) through the entire NMs with an inverted Zeiss Axiovert. We reconstructed the NMs using Axiovision or Zen blue (Zeiss software). At least two different blind observers counted the stained HCs/NM. We imaged a subset of larvae with a confocal microscope (Nikon Eclipse Ti-E Inverted Fluorescence Microscope) to document NMs.

### TUNEL staining

Prior to staining, we exposed previously anesthetized and fixed larvae to acetone for 7 minutes at −20°C, then rehydrated with 3 × 5min washes in PBST. Next, we digested larvae in Proteinase K (10 μg/mL) for 15 min and subsequently washed them 3 × 5min in PBST. For the subsequent steps we followed the *In-Situ* Cell Death Detection Kit, TMR Red (Roche #12156792910) directions adjusting the volumes as needed for whole larvae.

### Embryo and larval survival curve

Starting at 3dpf and for the following 4 days (until 7dpf), we raise larvae in one of the following media: SW, SW + Val_1_, or SW + VPA_150_ that was replaced daily. Each day, we checked thoroughly the health status namely by visually assessing under the stereoscope, the presence of morphological/developmental defects (heartbeat, blood flow spontaneous and provoked movement, presence of necrosis in internal organs, inflated swim bladder…) which can be easily done in larvae that are mostly transparent. For the survival count, we only retained thriving larvae devoid of any abnormality. We performed the same experiments on post copper-treated 5dpf larvae until 7dpf.

### Statistical Analysis

We averaged the number of HCs/NM in Graph Pad Prism 7 and assessed the statistical significance using simple T-tests in which we compared pairwise the experimental groups (Control vs. Val_1_) and (Control vs. VPA_150_). Survival curves were analyzed using non-parametric Kaplan-Meier estimator and significance was assessed using the Log-rank Mantel Cox test in Graph Pad Prism 7. *p* < 0.05 (*) was considered statistically significant and *p* < 0.001 (**) and above highly significant.

## Results

### Hair cell regeneration is impaired by Valerian crude root extracts

We used a simple HC regeneration assay schematically described (Figure 1a). Wildtype embryos were raised until 5dpf when we replaced the medium (system water, SW) with a Copper Sulfate II solution (Cu^++^, 10μM) for 2 hours (red bar and red arrow). This treatment triggered HC regeneration by synchronously killing all HCs in all NMs of the LL (Figure 1b, green dots) which we verified systematically, just after the treatment (+ 0hpt in c and d) by applying a fluorescent live fixable dye (FM1-43X, figure 1c, red). Next, we abundantly rinsed larvae before leaving them to recover for 24, 48 or 72hpt in one of the following media that was replaced daily: SW, SW + VPA_150_ (VPA_150_ = 150μM), or SW+Val_1_ (Val_1_= 1mg/ml). To determine the best concentrations to use for VPA and Val in our comparative approach, we settled for VPA_150_ based on the published highest well-tolerated concentration which was eliciting a strong decrease in HC regeneration (29, 30). We empirically determined that Val_1_ (= 1mg/ml) was the highest well-tolerated concentration for periods exceeding the duration of the regeneration assay (data not shown and survival curve in Figure 3). Representative NMs for each treatment and each time point (Figure 1c), displayed regenerated and physiologically active HCs (red, top panels and lower panels) at the respective time points (+0, + 24, + 48 and +72hpt) with all nuclei of the NMs counterstained with DAPI (blue, middle and lower panels). For each larva, we imaged at least 3 head and 3 trunk NMs (N = 3, n = 30 /treatment/hpt) and proceeded with 3D reconstruction in which we counted all stained HCs. Next, we averaged the number of HCs/NM and grafted them (Figure 1 d). At 24hpt, NMs from larvae recovering in SW had significantly more HCs (black bar ~ 4) than larvae in SW+Val_1_ (red bar ~2, p< 0.001), or in SW+VPA_150_ (gray bar, ~1, p< 0.0001). At 48hpt, larvae in SW had on average 6 HCs/NM, which was not significantly different from larvae in SW+Val_1_ with ~5 HCs/NM but strikingly different from larvae in SW+VPA_150_ with only ~2 HCs/NM. At 72hpt, larvae in SW had restored the number of HCs to a pre-copper treatment level (~8 HCs/NM) which was significantly more than in larvae treated with SW+Val_1_ (~5.5 HCs/NM, p <0.001), while NMs of SW+VPA_150_ treated larvae had at most 1 HC/NM (p< 0.00001). Taken together, we showed that treatments with crude extracts of Val_1_ were significantly hampering HC regeneration even if to a lesser extent than VPA_150_-treatments, which was not unexpected as the crude extracts most probably contained less active compound.

**Figure 1.**
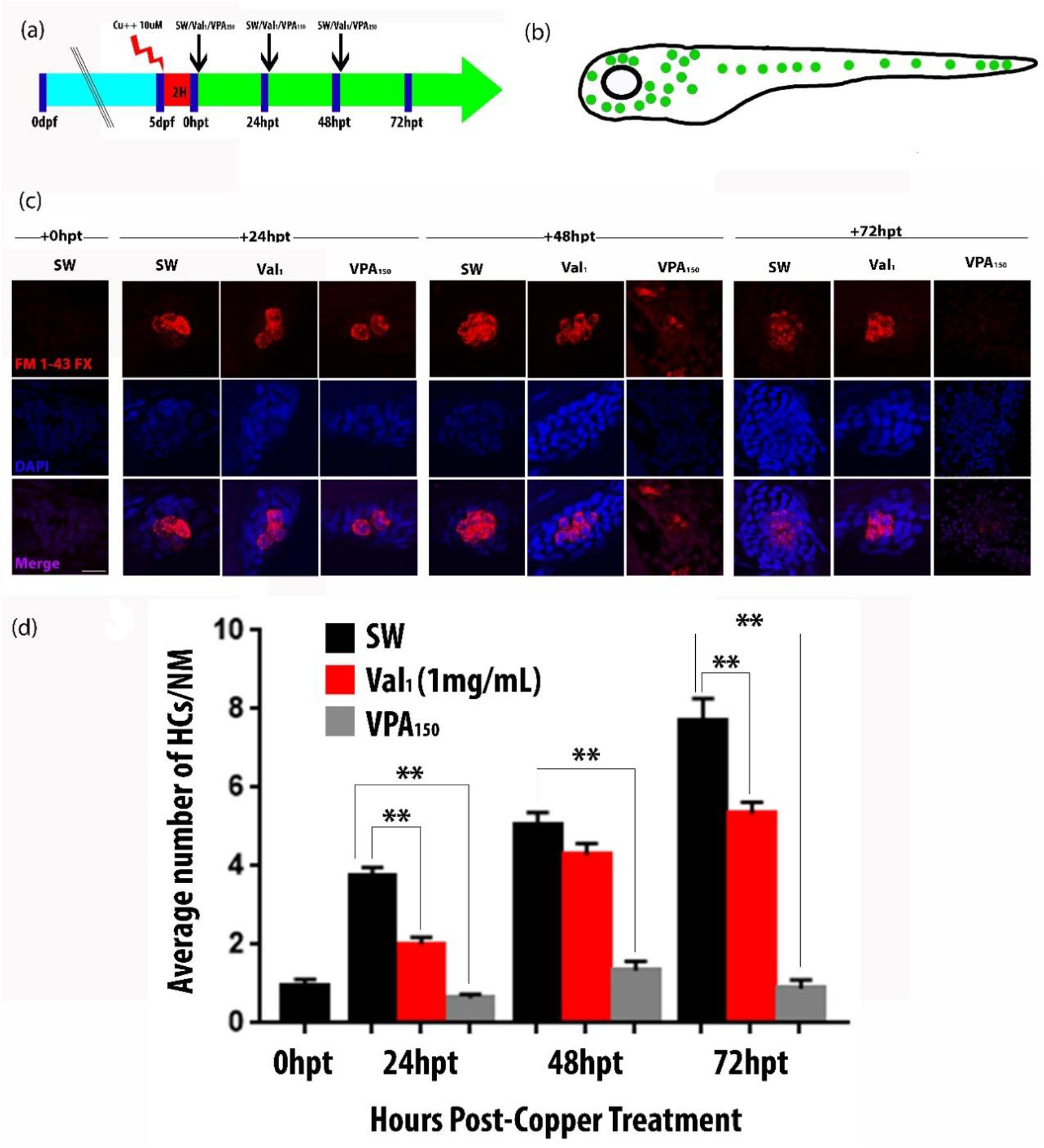
Hair cell (HC) regeneration in the larval lateral line (LL) is reduced after Val_1_ or VPA_150_ exposure. (a). Schematic of the experimental approach used for assessing HC-regeneration in the LL. Synchronized HC loss was triggered in 5dpf larvae by exposing them to 10μM Cu for 2h (red bar and arrow). Larvae were then left to recuperate for the following 3day post treatment (dpf) in either system water (SW), SW + Val_1_ (1mg/ml), or SW + VPA_150_ (150μM). HCs were counted at 4 different time points after copper treatment, namely at 0, 24, 48 and 72hour post-treatment (hpt) in all larvae (N = 3, n = 30/treatment/time point). (b). Schematic of a 5dpf larva showing the stereotypical distribution of neuromasts (NMs, green dots) in the head, trunk and tail composing the lateral line (LL). (c). Confocal images of representative NMs at the indicated observation stage after copper treatment (hpt). Physiologically active HCs are labelled with the live dye (FM1-43X, red upper and lower panels) and all nuclei in a NM are counter-stained with DAPI (blue, middle, and lower panels). (d). Graphic representation of the average number of HCs/NM at the indicated observation stage (hpt) after regenerating in SW (black bars), in SW + Val_1_ (red bars), or in SW + VPA_150_ (grey bars). Scale bar in c = 30μM. Errors bars represent s.e.m, and statistical significance is indicated as following: \*\**p* < 0.001; \*\*\**p* <0.0001.

### Valerian treatments do not affect HC survival

VPA_150_ treatments were previously shown to have no effect on HCs survival during regeneration (29, 30), suggesting that the decline in HC regeneration was apoptosis-independent. To verify if this was also the case for Val_1_-treatments, we performed a TUNEL assay that fluorescently end-labelled fragmented DNA, a hallmark of apoptotic cells. Furthermore, to identify which cells were dying we performed the experiment in the transgenic line (*Tg* (tnks1bp1: EGFP)) which is expressing GFP in support cells (SCs) but not HCs in all NMs (Figure 2, green). As expected, when we observed NMs just after the copper treatment (+0hpt) we found a significant number of TUNEL-labelled cells (red). Most of them were not expressing GFP presumably representing the remnant of dying HCs, and/or macrophages that were recruited to the site and had engulfed HC debris, as previously described (32). We also noted some GFP-expressing cells that were TUNNEL-labelled indicating that at least some SCs were affected by the copper treatment, as previously described (21). Strikingly, we found no NMs with TUNEL-positive cells in any subsequent regenerative stages (+24, +48 and +72hpt) for neither treatment (Figure 2a, SW: top panels, SW+Val_1_: bottom panels). We counted TUNEL-positive cells in all NMs at all stages (N = 2, and n = 20/stage), averaged and graphed the results (Figure 2b). At + 0hpt, we found, as expected, ~4 TUNEL positive cells/NM, but quasi none at subsequent stages for the rare exception of scattered labelling in a few NMs for both post-copper treatments, SW (black bars) and SW+Val_1_ (red bars), which were not significantly different from each other. Taken together, this was indicating that Val_1_-treatments were not killing HCs or SCs during HC-regeneration, thus suggesting that comparable to VPA_150_-treatments, the reduced HCs regeneration was apoptosis-independent and possibly via a similar mode of action.

**Figure 2.**
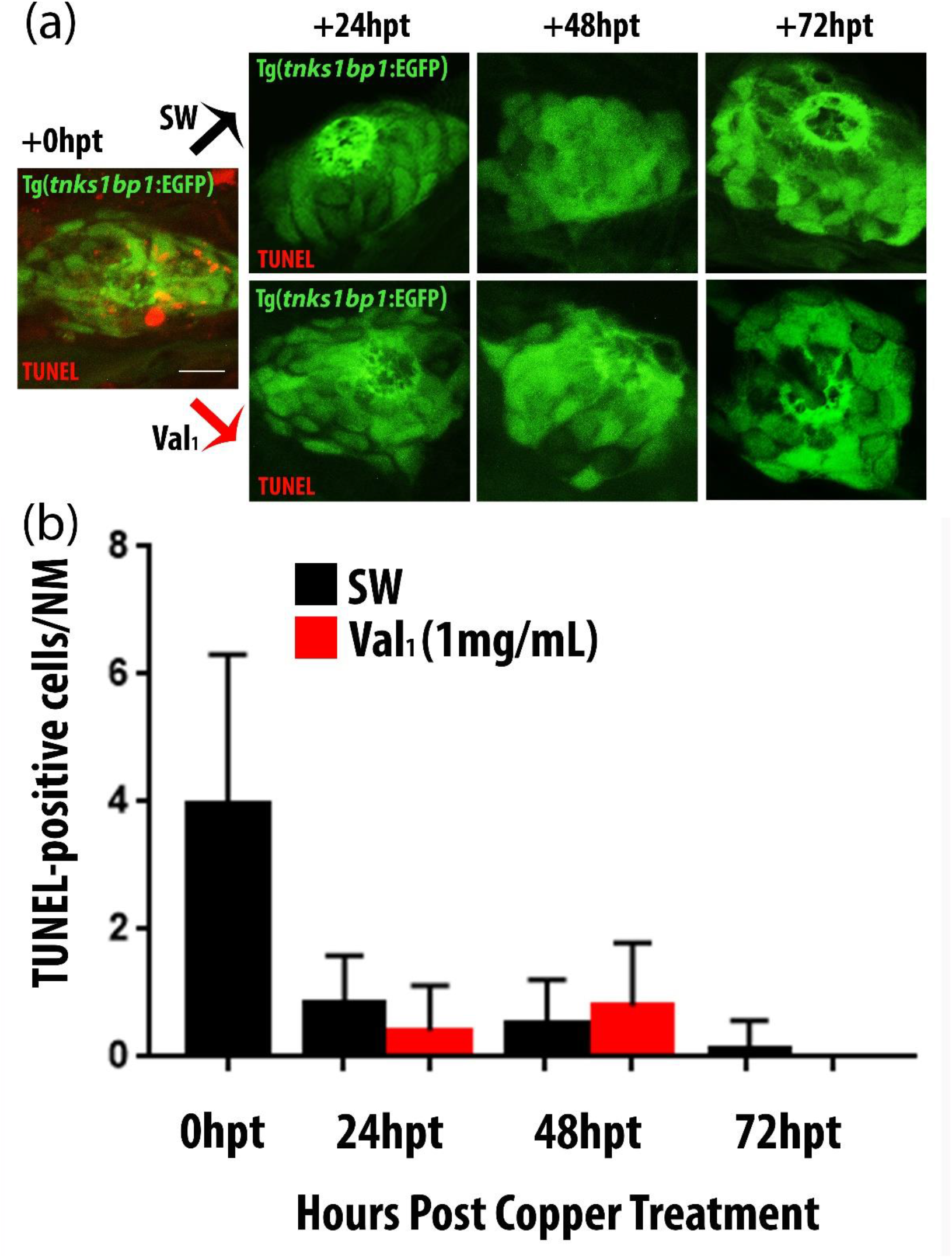
TUNEL-labelling of apoptotic cells in regenerating NMs after copper-treatment. (a). Confocal images of NMs of Tg (*tnks1bp1: EGFP*) larvae which are expressing GFP in all SCs (green) that were treated in a TUNEL assay end-labelling fragmented DNA (red). All animals were previously copper-treated to kill HCs and trigger HC-regeneration. Just after copper-treatment (0hpt, left panel) several apoptotic cells are visible. Larvae left to recover in SW (top panels) or in SW + Val_1_ (bottom panels) were TUNEL-probed and imaged at 24, 48 or 72hpt, but showed no apoptotic cells. (b). Average TUNEL-positive labelled cells/NM after the indicated recovery time post copper-treatment in SW (black bars), or SW + Val_1_ (red bars). Error bars represent the s.e.m, scale bar in a = 10 μM.

### Larval survival rate is not affected by Valerian root extracts treatments

Next, we wanted to exclude that decreased HC regeneration with VPA_150_ and Val_1_-treatments was due to general toxicity affecting proper development, general health, or survival of larvae. Thus, we treated larvae starting at 3dpf until 7dpf with similar concentrations (VPA_150_ and Val_1_) for periods exceeding the duration of the HC regeneration assay (from 5 to 7dpf). We found that none of the 1 to 4 day-long continuous treatments (N= 3, n= 22/treatment) significantly affected proper development, general health (assessed by several vital criteria: heartbeat, blood flow, spontaneous and provoked movements, and inflated swim bladder data not shown), or the survival rate (Figure 3a).

**Figure 3.**
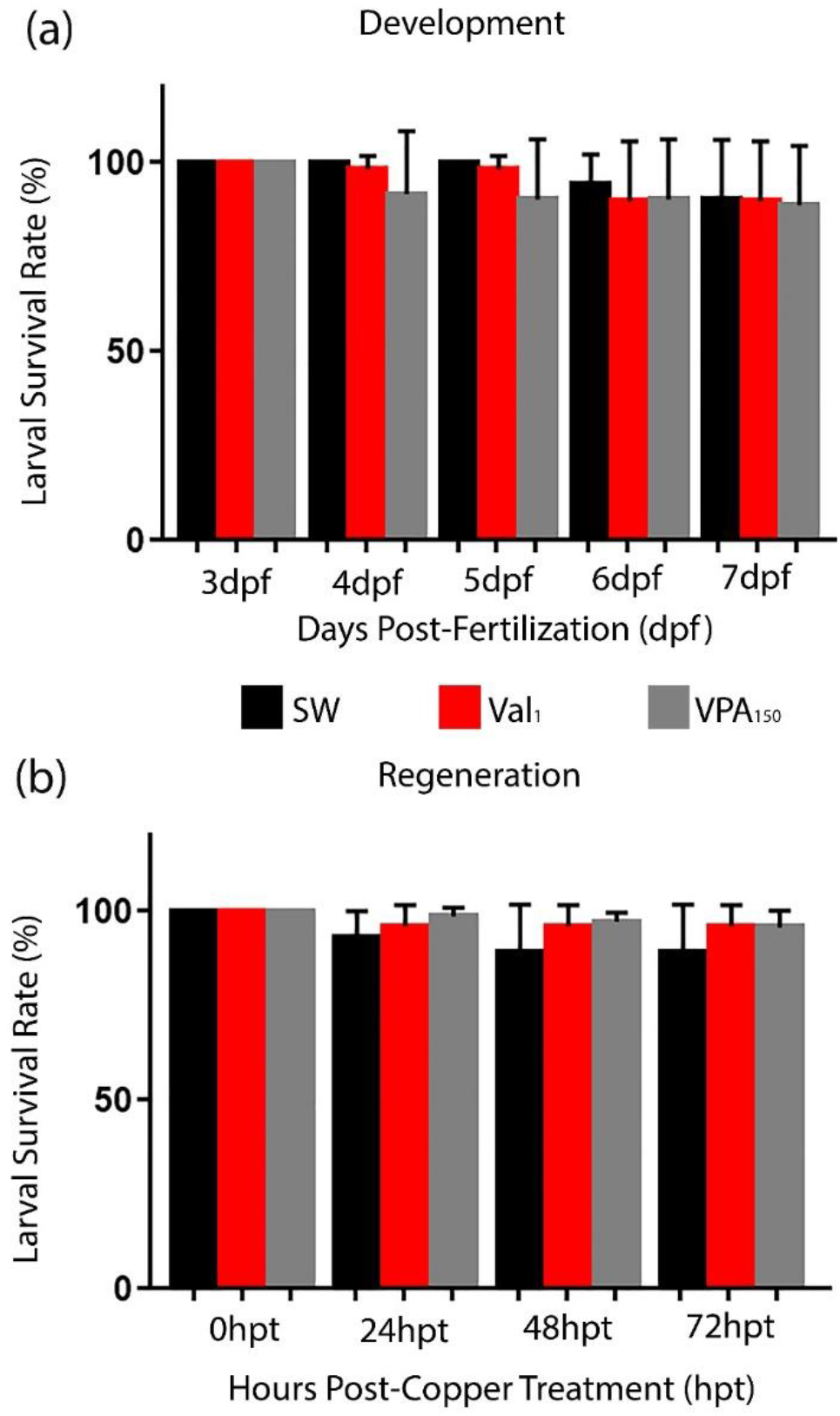
Survival of larvae after chronic exposure to Val_1_ or VPA_150_ with or without copper-treatment. (a). Percentage of survival rate of larvae raised from 3dpf onward until the indicated dpf in SW (black bar), SW+ Val_1_ (red bar) or in SW + VPA_150_ (grey bar) without copper-treatment. (b). Percentage of survival rate of larva after a punctual 2h copper-treatment at 5dpf followed by exposure to SW alone (black bar), to SW+ Val_1_ (red bar), or to SW + VPA_150_ (grey bar) for the following 3 days post copper-treatment (+24, + 48, and +72hpt). Errors bars represent the s.e.m.

Finally, to exclude the fact that there might be a compound effect of the VPA_150_ and Val_1_-treatments with the copper-treatment that we used to trigger HC-death and regeneration, we next closely monitored general health and survival of animals after exposure to copper and subsequent Val_1_ or VPA_150_-treatments (N=3, and n=25/treatments). Again, we found no overt health problem, or significant mortality (Figure 3b). Taken together, neither VPA_150_ nor Val_1_-treatments, in association with or without copper-treatment, had obvious detrimental consequences on general health or survival of the treated larvae, suggesting that the decreased HC regeneration was specific and not a secondary effect caused by treatment toxicity.

## Discussion and conclusions

We assessed the feasibility of using crude plant extracts of unknown composition which were made from powdered Valerian roots in a simple HC regeneration assay in the zebrafish larval lateral line (LL). We elaborated a central postulate on three facts: first, a major component present in any Val extract is Valeric acid (VA) which was previously demonstrated in this well studied plant (27).; second, valproic acid (= valproate, VPA) is the synthetic analog of VA (33); and third, VPA significantly reduces HC regeneration in the zebrafish LL (29, 30). Thus, we reasoned that we should be able to elicit an effect comparable to VPA when applying a crude extract of Val. This would serve as proof of principle that when an active ingredient is present in a non-purified mixture, it can elicit a readable and specific effect in a simple up-scalable HC regeneration assay. We showed that with this readily up-scalable method we could effectively detect the postulated reduction in HC regeneration when treating regenerating animals with a preparation of Val at 1mg/ml (Val_1_), the highest well-tolerated concentration that we had previously determined. Importantly, we also demonstrated that the reduced HC regeneration was not due to ototoxicity or overall toxicity, arguing that we were detecting a specific effect.

The larval zebrafish has been extensively used in drug discovery and toxicology screens (for review (34)), and so has the lateral line sensory organ (13–15, 17, 18, 35–37). Until now, purified molecules were screened often using pre-existing small molecule libraries, which when FDA-approved have the obvious advantage of offering a ready-to-go drug in case of success. However, this comes with the clear limitation that only already known, synthetic molecules will be interrogated. We propose that systematic screening of crude preparations could be set up after a careful selection based on the ethnobotanical documented regeneration potential of plant/fungal crude preparations, as described in the abundant literature and available databases (i.e http://herb.umd.umich.edu/, http://www.herbmed.org/, and http://www.leffingwell.com/plants.htm). Standardized screens would allow to rapidly interrogate any complex mixture for the potential of decreasing or enhancing HC-survival and regeneration. The same inexpensive and rapid screens could then be used in successive purification steps to isolate the active compound(s) in extracts of interest. We demonstrated in these proof-of-principal experiments, efficacity and safety of valerian crude root extracts of unknown composition in an *in vivo* HC regeneration assay. Our work, while extending the potential of an up-scalable approach, has obvious short-comings. Extracts can generate false positives because of potential synergistic effects of the hundreds of compounds present in a mixture which might mask the effect of any single ingredient. Extracts might not be testable at an effective concentration without becoming toxic. Likewise, false positives could be generated by synergistic effects that might be difficult to recreate with purified solutions. Nevertheless, using these cost-effective rapid screens could significantly increase discovery rate of new molecules by enabling the exploration of the vast NP pharmacopoeia and therefore promote translational solutions for hearing and balance disorders.

## Acknowledgements

We want to thank the following undergraduate students: Loranie Colom, Juan Cantres-Velez, Kenny Colón and Keisy Bonano as well as to the fish room technician Neidibel Martinez for help with husbandry and fish care. Thanks to Bismarck Madera, (Platform imaging technician), and to management for graciously for assisting with and letting us use the confocal facility in the Molecular Science Building (San Juan, PR)

## References

1. Wilson BS, Tucci DL, Merson MH, O’Donoghue GM. Global hearing health care: new findings and perspectives. Lancet. 2017;390(10111):2503–15.

2. Bommakanti K, Iyer JS, Stankovic KM. Cochlear histopathology in human genetic hearing loss: State of the science and future prospects. Hear Res. 2019;382:107785.

3. Muller U, Barr-Gillespie PG. New treatment options for hearing loss. Nat Rev Drug Discov. 2015;14(5):346–65.

4. Rubel EW, Furrer SA, Stone JS. A brief history of hair cell regeneration research and speculations on the future. Hear Res. 2013;297:42–51.

5. Corwin JT, Cotanche DA. Regeneration of sensory hair cells after acoustic trauma. Science. 1988;240(4860):1772–4.

6. Ryals BM, Rubel EW. Hair cell regeneration after acoustic trauma in adult Coturnix quail. Science. 1988;240(4860):1774–6.

7. Cox BC, Chai R, Lenoir A, Liu Z, Zhang L, Nguyen DH. et al. Spontaneous hair cell regeneration in the neonatal mouse cochlea in vivo. Development. 2014;141(4):816–29.

8. Hu L, Lu J, Chiang H, Wu H, Edge AS, Shi F. Diphtheria Toxin-Induced Cell Death Triggers Wnt-Dependent Hair Cell Regeneration in Neonatal Mice. J Neurosci. 2016;36(36):9479–89.

9. Burns JC, Stone JS. Development and regeneration of vestibular hair cells in mammals. Semin Cell Dev Biol. 2017;65:96–105.

10. Roccio M, Senn P, Heller S. Novel insights into inner ear development and regeneration for targeted hearing loss therapies. Hear Res. 2019:107859.

11. Ghysen A, Dambly-Chaudiere C. The lateral line microcosmos. Genes Dev. 2007;21(17):2118–30.

12. Ton C, Parng C. The use of zebrafish for assessing ototoxic and otoprotective agents. Hear Res. 2005;208(1-2):79–88.

13. Owens KN, Santos F, Roberts B, Linbo T, Coffin AB, Knisely AJ, et al. Identification of genetic and chemical modulators of zebrafish mechanosensory hair cell death. PLoS Genet. 2008;4(2):e1000020.

14. Chiu LL, Cunningham LL, Raible DW, Rubel EW, Ou HC. Using the zebrafish lateral line to screen for ototoxicity. J Assoc Res Otolaryngol. 2008;9(2):178–90.

15. Hirose Y, Simon JA, Ou HC. Hair cell toxicity in anti-cancer drugs: evaluating an anti-cancer drug library for independent and synergistic toxic effects on hair cells using the zebrafish lateral line. J Assoc Res Otolaryngol. 2011;12(6):719–28.

16. Thomas AJ, Wu P, Raible DW, Rubel EW, Simon JA, Ou HC. Identification of small molecule inhibitors of cisplatin-induced hair cell death: results of a 10,000 compound screen in the zebrafish lateral line. Otol Neurotol. 2015;36(3):519–25.

17. Ou HC, Cunningham LL, Francis SP, Brandon CS, Simon JA, Raible DW. et al. Identification of FDA-approved drugs and bioactives that protect hair cells in the zebrafish (Danio rerio) lateral line and mouse (Mus musculus) utricle. J Assoc Res Otolaryngol. 2009;10(2):191–203.

18. Namdaran P, Reinhart KE, Owens KN, Raible DW, Rubel EW. Identification of modulators of hair cell regeneration in the zebrafish lateral line. J Neurosci. 2012;32(10):3516–28.

19. Ratsch C. The Encyclopedia of psychoactive plants: Ethnopharmacology and its applications. Press PS, editor. US2005.

20. Behra M, Bradsher J, Sougrat R, Gallardo V, Allende ML, Burgess SM. Phoenix is required for mechanosensory hair cell regeneration in the zebrafish lateral line. PLoS Genet. 2009;5(4):e1000455.

21. Hernandez PP, Moreno V, Olivari FA, Allende ML. Sub-lethal concentrations of waterborne copper are toxic to lateral line neuromasts in zebrafish (Danio rerio). Hear Res. 2006;213(1-2):1–10.

22. Olivari FA, Hernandez PP, Allende ML. Acute copper exposure induces oxidative stress and cell death in lateral line hair cells of zebrafish larvae. Brain Res. 2008;1244:1–12.

23. Mackenzie SM, Raible DW. Proliferative regeneration of zebrafish lateral line hair cells after different ototoxic insults. PLoS One. 2012;7(10):e47257.

24. Pinto-Teixeira F, Viader-Llargues O, Torres-Mejia E, Turan M, Gonzalez-Gualda E, Pola-Morell L. et al. Inexhaustible hair-cell regeneration in young and aged zebrafish. Biol Open. 2015;4(7):903–9.

25. Romero-Carvajal A, Navajas Acedo J, Jiang L, Kozlovskaja-Gumbriene A, Alexander R, Li H. et al. Regeneration of Sensory Hair Cells Requires Localized Interactions between the Notch and Wnt Pathways. Dev Cell. 2015;34(3):267–82.

26. Ma EY, Rubel EW, Raible DW. Notch signaling regulates the extent of hair cell regeneration in the zebrafish lateral line. J Neurosci. 2008;28(9):2261–73.

27. Patočka J J, J. Biomedically relevant chemical constituents of Valeriana officinalis. Journal of Applied Biomedicine. 2010;8(1):11–8.

28. Chiu CT, Wang Z, Hunsberger JG, Chuang DM. Therapeutic potential of mood stabilizers lithium and valproic acid: beyond bipolar disorder. Pharmacol Rev. 2013;65(1):105–42.

29. He Y, Cai C, Tang D, Sun S, Li H. Effect of histone deacetylase inhibitors trichostatin A and valproic acid on hair cell regeneration in zebrafish lateral line neuromasts. Front Cell Neurosci. 2014;8:382.

30. He Y, Mei H, Yu H, Sun S, Ni W, Li H. Role of histone deacetylase activity in the developing lateral line neuromast of zebrafish larvae. Exp Mol Med. 2014;46:e94.

31. Westerfield M. The zebrafish book: a guide to the laboratory use of the zebrafish (Danio Rerio). Eugene: University of Oregon Press. 2000;4th edition.

32. Williams JA, Holder N. Cell turnover in neuromasts of zebrafish larvae. Hear Res. 2000;143(1-2):171–81.

33. Henry TR. The history of valproate in clinical neuroscience. Psychopharmacol Bull. 2003;37 Suppl 2:5–16.

34. Cassar S, Adatto I, Freeman JL, Gamse JT, Iturria I, Lawrence C, et al. Use of Zebrafish in Drug Discovery Toxicology. Chem Res Toxicol. 2019.

35. Froehlicher M, Liedtke A, Groh KJ, Neuhauss SC, Segner H, Eggen RI. Zebrafish (Danio rerio) neuromast: promising biological endpoint linking developmental and toxicological studies. Aquat Toxicol. 2009;95(4):307–19.

36. Ou H, Simon JA, Rubel EW, Raible DW. Screening for chemicals that affect hair cell death and survival in the zebrafish lateral line. Hear Res. 2012;288(1-2):58–66.

37. Ou HC, Santos F, Raible DW, Simon JA, Rubel EW. Drug screening for hearing loss: using the zebrafish lateral line to screen for drugs that prevent and cause hearing loss. Drug Discov Today. 2010;15(7-8):265–71.

